# On convolutional neural networks for selection inference: revealing the lurking role of preprocessing, and the surprising effectiveness of summary statistics

**DOI:** 10.1101/2023.02.26.530156

**Authors:** Ryan M Cecil, Lauren A Sugden

**Affiliations:** Department of Mathematics and Computer Science, Duquesne University, Pittsburgh, PA, USA; Department of Statistics, University of Pittsburgh, Pittsburgh, PA, USA

## Abstract

A central challenge in population genetics is the detection of genomic footprints of selection. As machine learning tools including convolutional neural networks (CNNs) have become more sophisticated and applied more broadly, these provide a logical next step for increasing our power to learn and detect such patterns; indeed, CNNs trained on simulated genome sequences have recently been shown to be highly effective at this task. Unlike previous approaches, which rely upon human-crafted summary statistics, these methods are able to be applied directly to raw genomic data, allowing them to potentially learn new signatures that, if well-understood, could improve the current theory surrounding selective sweeps. Towards this end, we examine a representative CNN from the literature, paring it down to the minimal complexity needed to maintain comparable performance; this low-complexity CNN allows us to directly interpret the learned evolutionary signatures. We then validate these patterns in more complex models using metrics that evaluate feature importance. Our findings reveal that common preprocessing steps play a central role in the learned prediction method, most commonly resulting in models that mimic a previously-defined summary statistic, which itself achieves similarly high accuracy. In other cases, preprocessing steps introduce artifacts that can lead to “shortcut learning”. We conclude that human decisions still wield significant influence on these methods, hindering their potential to learn novel signatures. To gain new insights into the workings of evolutionary processes through the use of machine learning, we propose that the field focus on methods that avoid human-dependent preprocessing.

**Author summary:** The ever-increasing power and complexity of machine learning tools presents the scientific community with both unique opportunities and unique challenges. On the one hand, these data-driven approaches have led to state-of-the-art advances on a variety of research problems spanning many fields. On the other, these apparent performance improvements come at the cost of interpretability: it is difficult to know how the model makes its predictions. This is compounded by the computational sophistication of machine learning models which can lend a deceptive air of objectivity, often masking ways in which human bias may be baked into the modeling decisions or the data itself. We present here a case study, examining these issues in the context of a central problem in population genetics: detecting patterns of selection from genome data. Through this application, we show how human decision-making can influence model predictions behind the scenes, sometimes encouraging the model to see what we want it to see, and at other times, presenting the model with signals that allow it to circumvent the learning process. By understanding how these models work, and how they fail, we have a chance of creating new frameworks that are more robust to human biases.

## Introduction

Over the last 30 years, methods for detecting positive selection from present-day genome data have become more sophisticated and more powerful. Early summary statistics were designed to detect the signature of strong selection, or a “selective sweep” based on its effect on the site frequency spectrum, relying on a signature of over-abundance of low- and high-frequency derived alleles [48, 11]. Later, summary statistics were developed that capitalized on the signature of extended haplotype homozygosity (EHH)[40, 51, 15, 41, 12], a result of adaptive mutations rising quickly in frequency along with neighboring “hitchhiking” alleles. The 2010s saw the development of composite classification models designed to increase the power to detect sweeps by combining multiple summary statistics in various supervised classification frameworks, training these classifiers on simulated genomic data[42, 20, 47, 45, 30, 37].

More recently, advances in machine learning have presented an exciting prospect: these models might be able to learn patterns of natural selection directly from raw data, thereby extracting more information than is available through current handcrafted summary statistics. For the purposes of detecting natural selection, multiple deep learning architectures have recently been proposed that can detect signatures of natural selection from sequence data with high accuracy[43]. Open questions remain, however, concerning the interpretability of these methods and how much power they add in various circumstances beyond existing methods.

While deep learning approaches have become state-of-the-art for a variety of tasks such as image processing [29, 28, 22, 26], natural language processing [53, 7, 1], game playing [36, 46], and protein folding [44, 24], these models are often highly complex and not well understood. A few consequences of this are worth consideration. First, there are infinitely many possible model architectures, and identifying the optimal architecture for a given problem can be difficult. Second, once an architecture is chosen, it can be difficult to explain the resulting model’s predictions. Third, deep learning models can be particularly susceptible to over-fitting on artifacts, thereby performing “shortcut learning,” in which the model learns decision rules that perform well on a training set, but do not generalize to real-world datasets[16].

The recently proposed state-of-the-art deep learning approaches for detecting selective sweeps are convolutional neural networks (CNNs) [14, 49, 9, 6]. These approaches vary widely in their choice of architecture, including the number of layers, the number and size of convolution kernels, and methods of regularization (see S1 File). While all methods report high accuracy on testing data, what is as yet unexplored is how these models work, what artifacts they might be susceptible to, and what insights they might be able to provide to our current theory. In an ideal case, by understanding what additional information is deemed valuable by these complex models, it could be possible to improve the design of easy-to-use summary statistics beyond their current capabilities.

To address these questions, we first develop a very low-complexity CNN that we name “mini-CNN,” which maintains high performance while allowing for more transparency into the classifier’s inner workings. In addition, to interpret the more complex models, we draw on recent advances from the field of explainable AI [31, 38, 39], in particular using post-hoc interpretability methods built for visualizing and explaining feature importance for predictions after model training. Using these tools, we show empirically that typical preprocessing pipelines induce the CNN models for sweep detection to closely mimic handcrafted summary statistics; their decision rules are particularly similar to that of Garud’s H1, which typically performs at least as well at this classification task. We also explore how some preprocessing artifacts can lead to shortcut learning, resulting in models that would not be expected to generalize well.

## Results

### CNNs for Detecting Selection

Multiple CNN methods have been proposed to solve tasks within the field of population genetic inference, with each following a common approach. First, training data is generated through a demographic model of choice using simulation software. Each sample, consisting of a set of haplotypes, is then organized into an image where rows represent individual haplotypes and columnns represent genomic sites. These images undergo processing, and are standardized to a uniform width using varying techniques. Then, a CNN architecture is chosen and the model is trained and tested on the simulated genomic image data. Afterwards, the model may be applied to genome data from real populations. Figure 1a illustrates a general example of the process of applying a CNN to sampled haplotype data.

**Fig 1.**
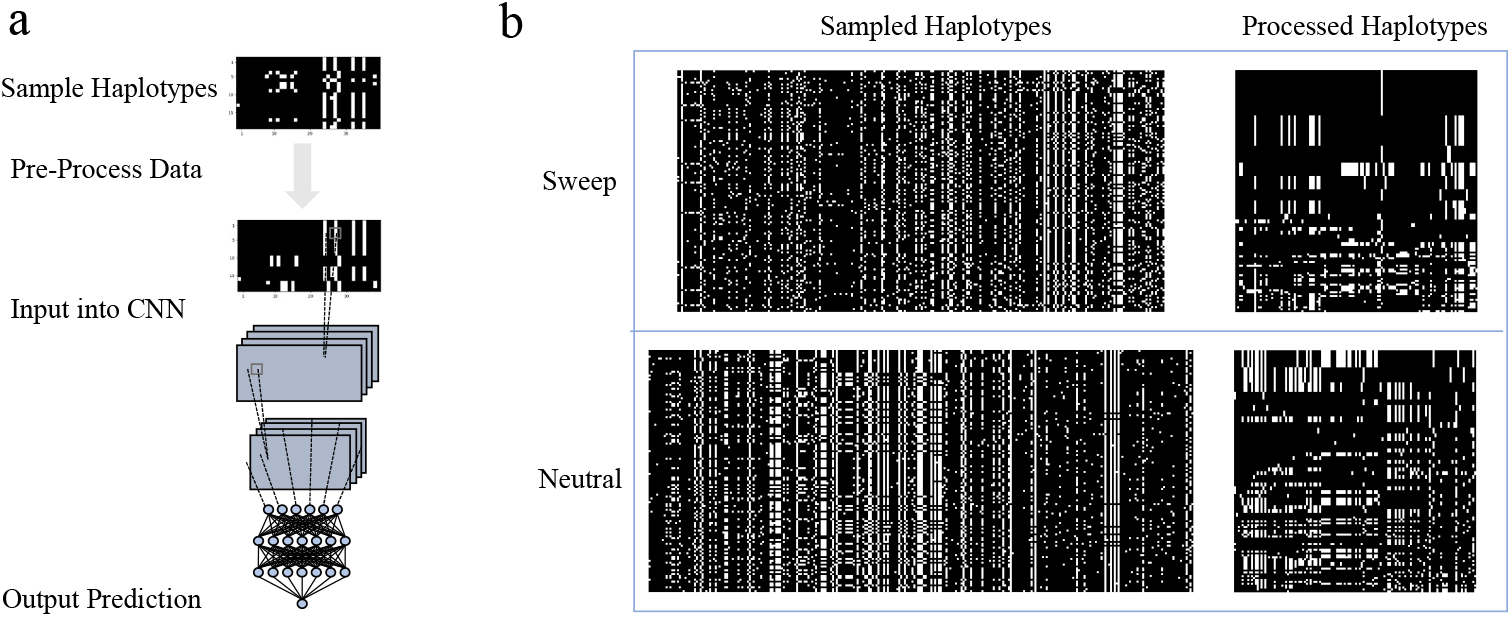
Typical workflow for CNNs for sweep detection. a: Raw sequence data is represented as an image with rows corresponding to sampled haplotypes and columns corresponding to variable loci. Pre-processing steps ensure that images are of uniform size, as required for the CNN, and can also rearrange rows and columns to bring out useful features for the CNN. The CNN itself can comprise a number of convolution, activation, and pooling layers, followed by a number of fully connected “dense” layers, ultimately resulting in a single numerical output representing a prediction (neutral or sweep). b: Examples of images representing raw data (left) and pre-processed data (right). In this case, we follow the conventions of Torada *et al*.[49], where the images are converted to major/minor polarization and filtered by allele frequency, the rows are sorted by genetic similarity, bringing the most common haplotype to the top, then the image undergoes width standardization.

S1 File contains an overview of recently proposed CNN methods, including their simulated data, preprocessing pipelines, and network architectures. Within the preprocessing pipeline, there exist three main approaches to generating images of fixed width: using an image resizing algorithm, sampling haplotypes of fixed width (either by including invariant sites, or allowing for varying genomic ranges), and zero-padding (see Methods). Notably, either before or after this resizing occurs, almost all of the methods [14, 49, 6] sort the haplotypes of the image by genetic similarity to create visible block-like features within the genetic image, with a block representing the most common haplotype at the top (see Figure 1b). In general, this has been shown to improve the performance of the model [14, 49]. Interestingly, the choice of CNN architecture varies widely across these works. The number of kernel dimensions, kernels, and convolutional layers alone range from 1D-2D, 4-256, and 1-4, respectively. They also vary in terms of the chosen demographic model and simulation software; altogether this makes direct comparison between the approaches tricky. A close look into how these CNN models make their predictions would aid in this regard.

To take a closer look into this class of CNN methods, we chose to implement and analyze the Imagene [49] model as a representative example, employing 2-dimensional convolutional kernels, and representing neither the least nor the most complex architecture. Like the other approaches, it sorts rows based on genetic similarity during preprocessing.

### Construction of mini-CNN and interpretation of its output

Interpretation of CNNs with complex architectures is notoriously difficult, so we first attempted to construct a minimal CNN that maintained high performance at the task of distinguishing between sweep and neutral simulations. Beginning with the Imagene model[49] as a representative example of CNNs for sweep detection, S1 Table shows the accuracy values as we reduce and remove the convolution, dense, ReLU, and max pooling layers for simulations of a single-population demographic model with a selection coefficient of *s* = 0.01. We find that the model with a single 2×1 kernel, followed by ReLU activation, then a single unit dense and sigmoid activation layer, maintains high performance (a difference of .3% accuracy to Imagene). We call this model mini-CNN (see S1 Fig for visualization of this model). Further simplification, such as removing the ReLU activation, results in a 18.4% accuracy loss.

Because of its simplicity, mini-CNN lends itself more easily to visual interpretation. In Figure 2a, we show the kernel weights for the 2×1 kernel, along with the dense weights map, which helps visualize the importance of different regions of the input image. The 2×1 kernel detects differences between two consecutive rows at a given locus, while the dark band at the top of the dense weights map, representing negative weights, indicates that the model interprets row-to-row differences near the top of the image as evidence against a sweep. Figure 2b illustrates how the model acts on an example sweep image. Because of row-sorting by genetic similarity, the large haplotype block at the top of the image does not contain row-to-row differences, leading to many 0’s (black pixels) at the top of the image after the 2×1 kernel convolution and ReLU activation. While 1’s (white pixels) in these positions would provide evidence for neutrality, these 0’s point the model in the opposite direction after the dense weights are applied.

**Fig 2.**
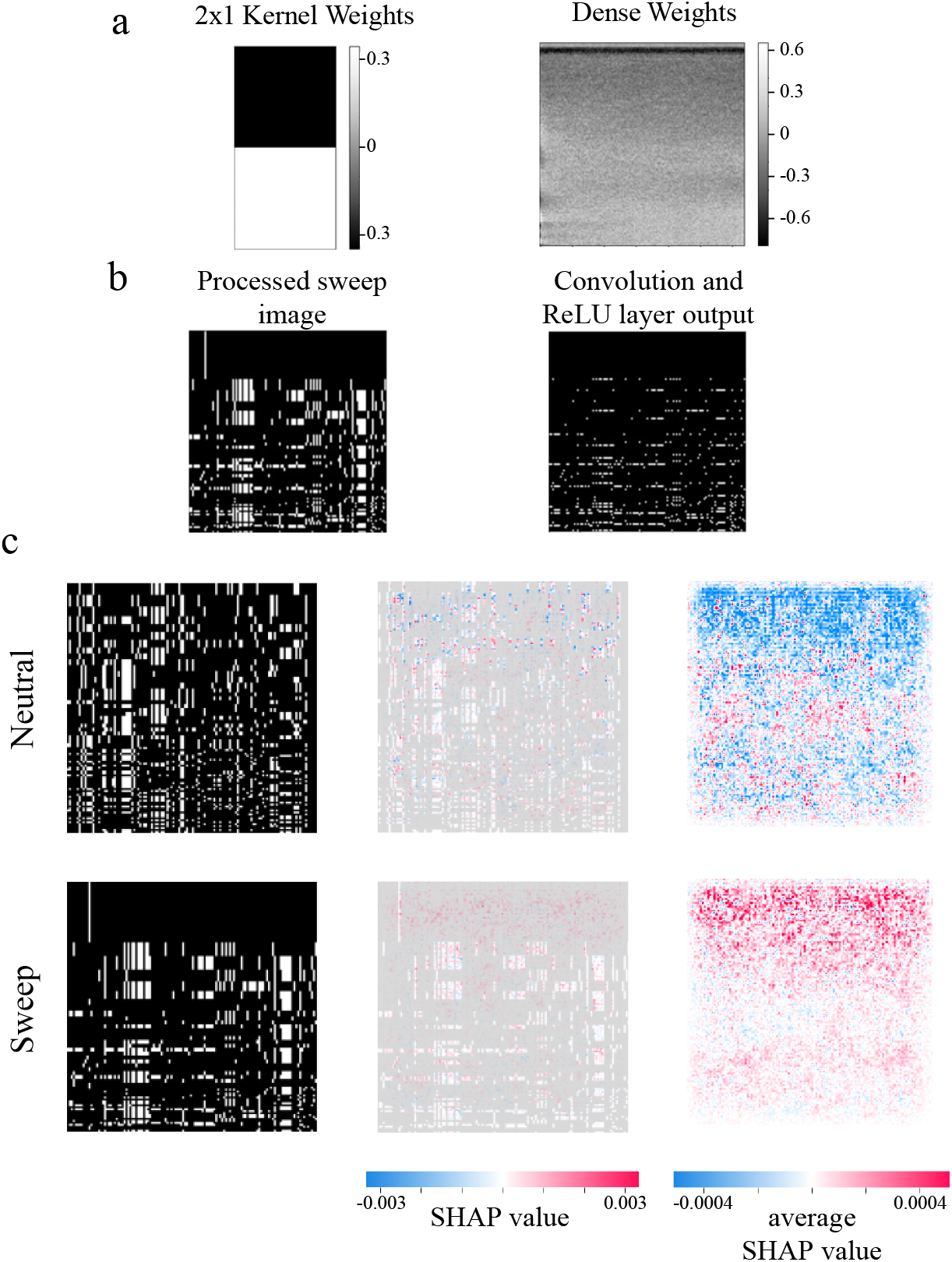
Opening the black box: visual explanations of mini-CNN and Imagene. a: Trained parameter values for mini-CNN. The 2×1 kernel will detect differences between consecutive rows. The dense weights map illustrates the linear weights that are applied to the output of the convolution and ReLU layer (depicted in panel b). The black band at the top of the weights indicates that the model is more likely to predict the image as neutral if there exists variation among the top rows. b: Example of a pre-processed image and its corresponding output after the convolution layer and ReLU activation. c: Visualization of Imagene with SHAP explanations. From left to right are examples of neutral and sweep processed images, SHAP values for the two image examples, and average SHAP values across 1000 neutral and sweep images. A negative SHAP value (blue) indicates that the pixel of interest contributes toward a prediction of neutral, while a positive SHAP value (red) indicates that the pixel of interest contributes toward a prediction of sweep. Similar to the black bar in the dense weights map in panel a, the large SHAP values located in the top region of the average Shap images indicate that Imagene focuses on the top block of the image to make its prediction.

Examining whether this pattern holds with more complex models requires alternative approaches; to this end, we applied an explanatory tool called SHAP [32] to the full Imagene model. SHAP values are a measure of the change in model prediction conditioned on the particular features, allowing us to see which pixels are most influential. Figure 2c shows SHAP values for each pixel within two example training images – one neutral, and one sweep – as well as what these SHAP values look like on average across 1000 neutral and 1000 sweep images. We see a similar pattern here, in which the regions that contribute the most to the model prediction appear near the top of the image, coinciding with the largest haplotype blocks.

### Comparison to Summary Statistics

The pattern we observe in which the CNN models pay the most attention to the rows at the top of the row-sorted image is not surprising; this coincides with existing theory about the signature left behind by a selective sweep; in particular, a sweep induces long shared haplotype blocks[40]. This is the basis of many current hand-crafted summary statistics, including iHS[51], Garud’s H statistics[15], nSL[13], and XP-EHH[41], with the last statistic requiring a reference population.

In Figure 3a, we measured the Spearman rank correlation between Imagene and mini-CNN output, and that of two commonly-used haplotype summary statistics, iHS and Garud’s H1. We found that Garud’s H1 in particular was highly correlated with both Imagene and mini-CNN. We also examined the correlation between the binary classification of each pair of methods (i.e. the proportion of simulations on which the classifiers agree; see Figure 3b). Here we find that Garud’s H1 agrees with both Imagene and mini-CNN 97% of the time in the single-population demographic model, and 89% of the time in the three-population demographic model.

**Fig 3.**
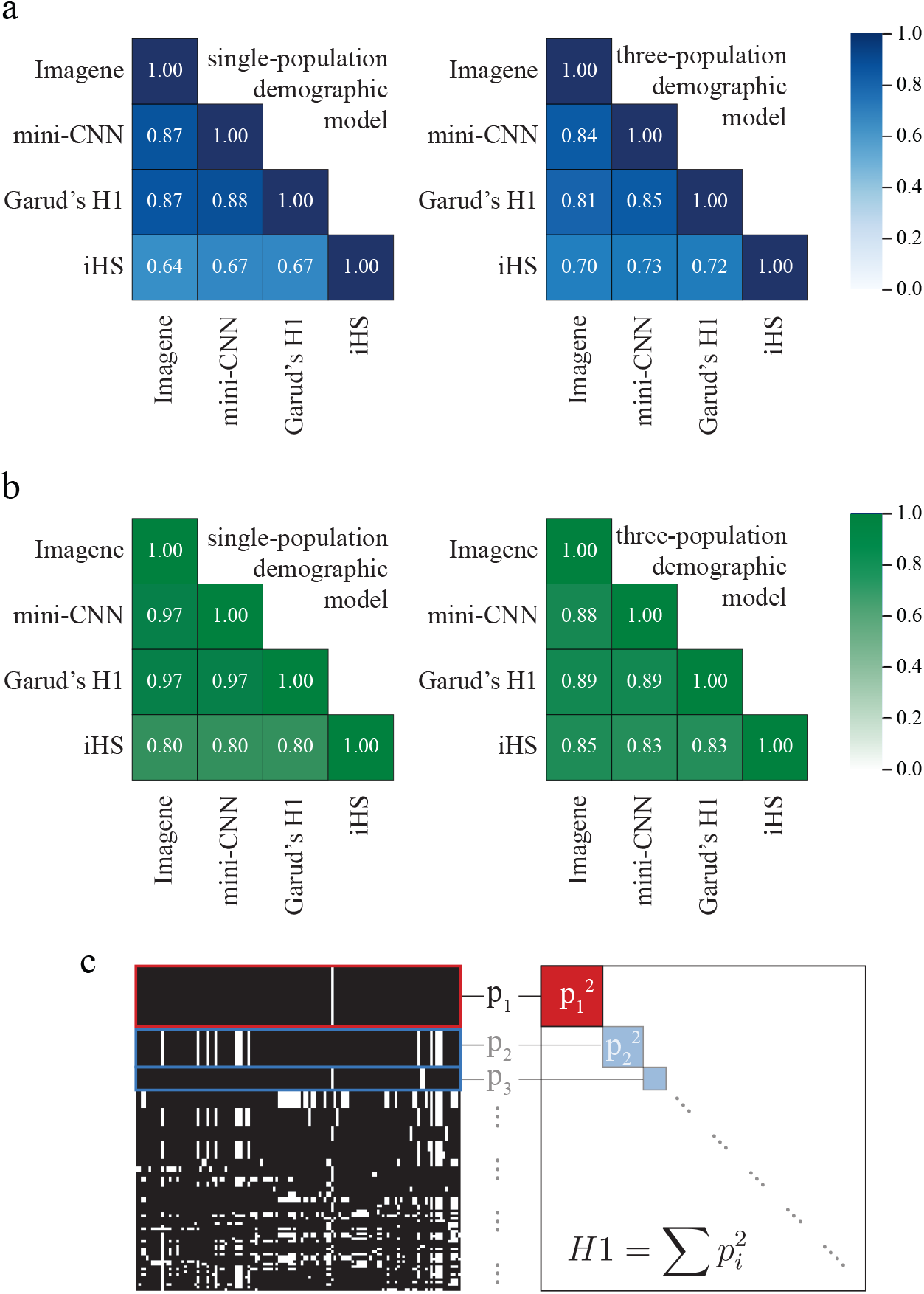
Comparison of CNNs with Garud’s H1 and iHS. a: Spearman rank correlation matrices comparing continuous output of CNN approaches and two summary statistics (Garud’s H1 and iHS) that measure haplotype homozygosity. Left: single-population demographic model simulated with msms. Right: three-population demographic model[19] with sweep in CEU. b: Correlation matrices comparing binary output of CNN appraches and two summary statistics. Left: single-population demographic model simulated with msms. Right: three-population demographic model[19] with sweep in CEU. Numbers shown are the proportion of simulations for which the two methods return the same classification (sweep or neutral), using optimal thresholds based on a training set. c: An example of a pre-processed image with row-sorting by genetic similarity (left). The haplotype with the highest frequency is brought to the top of the image. Calling this haplotype frequency *p*_1_, and the remaining haplotype frequencies *p*_2_, *p*_3_, …, Garud’s H1 statistic computes the sum of squared haplotype frequencies,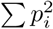. Both the CNNs and Garud’s H1 pick up on the signature of a high-frequency haplotype.

Figure 3c illustrates the design of Garud’s H1, alongside a training image pre-processed with genetic-similarity-based row sorting common to CNNs for sweep detection. Garud’s H1 operates by identifying shared haplotype blocks in a region, and computing the sum of squared haplotype frequencies. In the context of a pre-processed image then, Garud’s H1 is sensitive to the number of identical rows present at the top of the image, raising the possibility that in the process of row-sorting, the CNN is being set up to mimic the behavior of Garud’s H1.

In Table 1, we compare the accuracy of the CNN approaches and Garud’s H1 for the single-population demographic model. For two selection coefficients representing low and moderate selection strength, s=0.01 and s=0.005, Garud’s H1 actually *outperforms* both Imagene and mini-CNN.

**Table 1.**
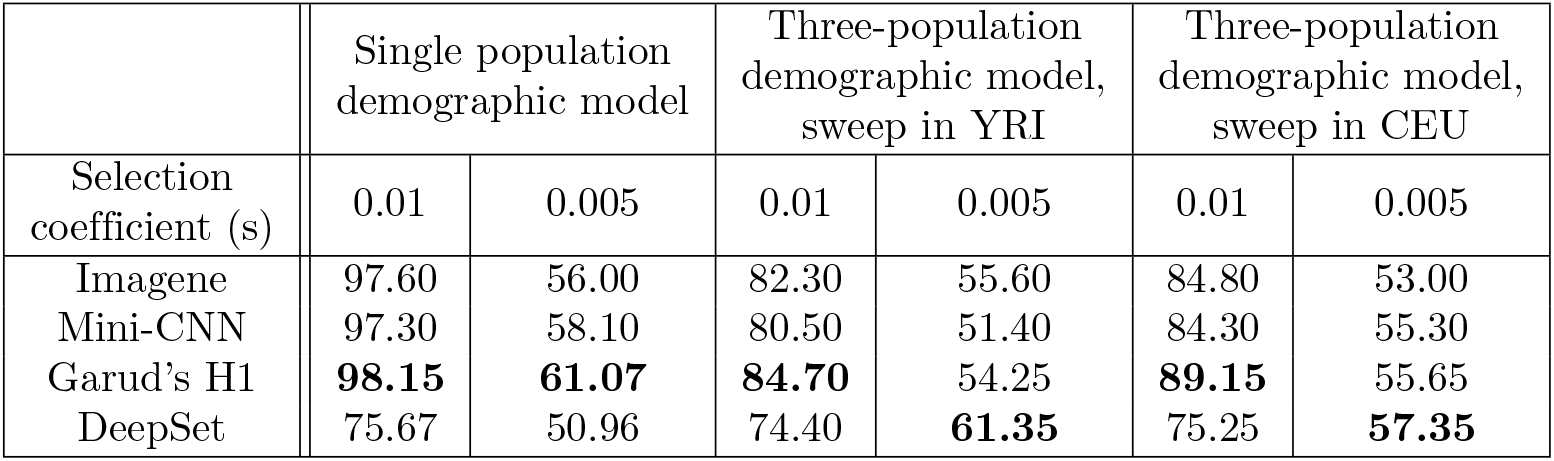
Performance Comparison. Performance values of multiple methods that were trained and tested on the same demographic model and selection coefficient. The performance values were found by computing the accuracy of the trained model on a balanced, held-out set.

### Performance in simulated human populations

A potential explanation for the performance of Garud’s H1 relative to the CNNs, and the performance of mini-CNN relative to Imagene, is that the single-population demographic model simulated with `msms` results in sequence data that is less noisy than genuine human genome data. This could produce selective sweep signals that are in effect too easy to detect, allowing for less complex models to perform on par with those that are more complex. Indeed, published CNNs for sweep detection (S1 File) have trained and tested their models on a variety of simulated demographic models, allowing for the possibility that these more complex models are necessary in more realistic human contexts.

To test these models on more realistic data, we used simulation software `slim` to simulate data under the three-population demographic model of Gravel *et al*.[19] which models the joint history of 1000 Genomes YRI, CEU, and CHB+JPT populations. This demographic model includes population bottlenecks and exponential growth, as well as migration among all three populations in the last 50,000 years. In S4 Fig, we show that the simulated neutral data replicate the site frequency spectrum of real 1000 Genome data[5] much more closely than the neutral simulations from the single-population model generated with `msms`. We also simulate two selection coefficients as above: *s* = 0.01 and *s* = 0.005, to investigate the compounded effect of more subtle sweeps.

Table 1 displays the accuracy of Imagene, mini-CNN, and Garud’s H1 for the three-population demographic model with sweeps in YRI and in CEU, for the two selection coefficients. As is expected due to the more complex demography, the performance of all three methods decline relative to their performance on the single-population demographic model. For stronger sweeps (*s* = 0.01), mini-CNN maintains a close performance to Imagene (a difference of 1.8% accuracy in the worse case), and for weaker sweeps (*s* = 0.005), mini-CNN’s performance is 4.2% lower than that of Imagene when the sweep is in YRI, and 2.3% higher when the sweep is in CEU. S1 Table contains accuracy values under this demographic model for different architectures representing the multiple levels of complexity between mini-CNN and Imagene.

Notably, the patterns we observed previously regarding Garud’s H1 hold in this context as well. Garud’s H1 outperforms both Imagene and mini-CNN on stronger sweeps, and performs on par with both on weaker sweeps. We note that all three methods seems to be near the edge of their detection range for these weaker sweeps (accuracy approaching 50%). In Figure 3a, we see that similar to the single-population demography case, Garud’s H1 is highly correlated with both Imagene and mini-CNN, and in S5 Fig and S7 Fig, we show that the dense weights map (in the case of mini-CNN) and SHAP values (in the case of Imagene) display the same trends that are shown in Figure 2: the models are most attentive to the haplotype blocks near the top of the image.

### CNN approaches are vulnerable to artifacts introduced in preprocessing

Following Torada *et al*.[49], we used an image resizing algorithm to attain genetic images of fixed width, but other methods have been proposed for this (see S1 File). To test whether the relationship between Garud’s H1 and the CNN models holds across other standardization methods, we implemented a form of zero padding, similar to the approach used in Flagel *et al*.[14]. In Figure 4a, we show examples of zero-padded sweep and neutral images, which display a common trend: because sweeps characteristically result in reduced haplotype diversity, an unprocessed sweep image typically contains fewer columns than an unprocessed neutral image, thus requiring more padding to attain a particular fixed width. In S1 Table, across the various simulation types, the accuracy values for the CNN models paired with image-width standardization via zero-padding are reported. Notably, when zero padding is applied, the performance of the CNN methods surpass that of the Garud’s H1, with the greatest improvements occurring for subtler sweeps. Further analysis, however, suggests that rather than revealing a domain in which the CNNs provide genuine insight beyond the capability of summary statistics, this increase in performance reflects “shortcut learning,” whereby models utilize artifacts in the data to improve predictive performance rather than meaningful signals.

**Fig 4.**
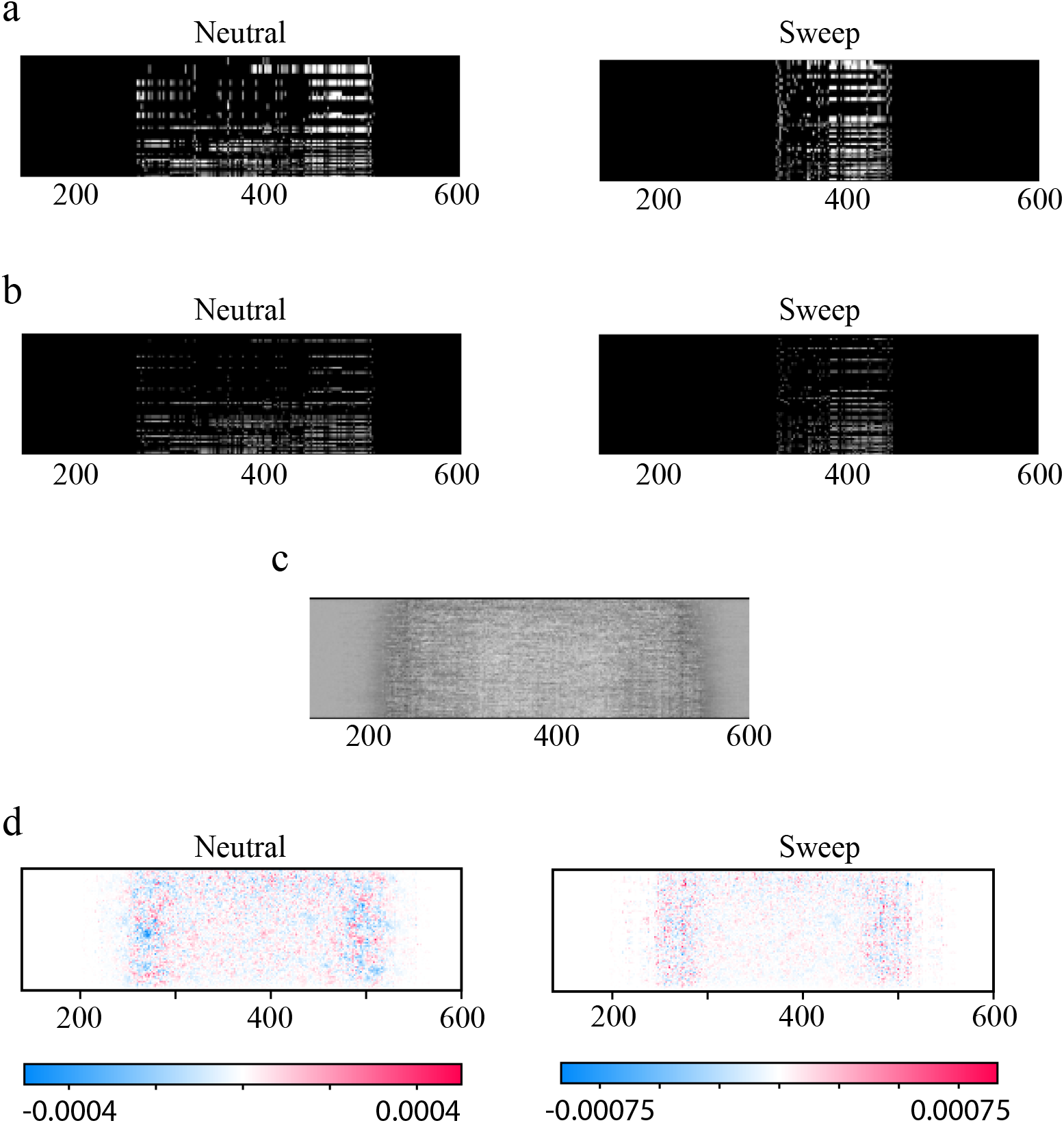
Image pre-processing has the potential to introduce artifacts: examining zero-padding in the presence of a low selection coefficient. a: Images processed with zero-padding from the three-population demographic model with a sweep in CEU with s=0.005. A signature of a selective sweep is a reduction in heterozygosity across the population; this manifests as fewer columns in the raw image after filtering for variable sites. Padding the image with columns of zeros thus results in larger blocks on either side of a sweep image relative to a neutral image. b: Output of the convolution and ReLU activation of mini-CNN for the images in panel a showing that the large blocks of zeros on either side of the image persist. c: Trained dense weights map for mini-CNN for the simulations in panel a show that the model is attentive to the width of the padding. d: SHAP explanations for Imagene on the same simulations likewise show greater importance along the side edges of the pre-processed image. Note that all images have been cropped for ease of visualization; in all cases, columns outside of the cropped region contain no information. Full images can be found in S9 Fig.

The effects of zero-padding, and potential artifacts introduced by its use, are shown in Figure 4b-c, which contain visualizations for mini-CNN and Imagene models trained with zero-padding on the three-population demographic model with a sweep in YRI. Interestingly, in Figure 4b, we see that in contrast to the dense weights map from figure 2a, the negative weights (darker regions) lie along the columns of the map at two locations on the left and right. This indicates that mini-CNN is searching for variation at these locations as evidence against a sweep. Since there exists no variation within the zero padded columns, mini-CNN is likely to predict a sweep when an image contains a large amount of zero padding, which in turn is dependent upon the number of columns of the unprocessed genetic image. Similar trends can be seen in the Imagene model with zero-padding. Figure 4b shows the output of the convolution and ReLU activation layers, which retain the original zero padding of the input. In Figure 4c, we show average SHAP values across 1000 neutral and 1000 sweep simulations for the Imagene model. Similar to above, the regions that contribute most to the model prediction lie along the columns of the left and right side.

Based on these results, it appears that image-width standardization via zero padding has the potential to create artifacts which allow the CNN model to observe the original width of the unprocessed simulated genetic image. While this enables the models to perform well in simulated data, this signature would not be expected to generalize to genuine human genome data, due to the vast range of recombination[27] and mutation[2] rates across the genome as a function of sequence context and other genome features; this degree of heterogeneity is not generally incorporated into simulation schemes.

To test the amount of information actually contained within this artifact, we crafted a summary statistic “Ncols” for detecting a sweep that simply counts the number of columns of the unprocessed image. In S1 Table, we show the accuracy of this method in comparison to the CNN models with zero padding. The accuracy values reveal that for the demographic models in which zero-padding improves performance the most – subtler sweeps in the three-population demographic model – this near-trivial summary statistic outperforms Imagene and mini-CNN regardless of the type of image-width standardization, and it also outperforms all other summary statistics including Garud’s H1.

We note that this artifact would not appear if image columns represented all genome positions (see e.g. [9]) rather than the collection of segregating sites, or the collection of segregating sites passing a minor allele frequency threshold. This has the downside, however, of introducing a large number of uninformative columns, thereby requiring wider images to accommodate the same number of segregating sites, and making the haplotype structure within the data less prominent as a visual signal.

### Comparison to Permutation Invariant Network

The learned prediction methods of CNNs built for the detection of selective sweeps appear to critically depend on human-designed methods of preprocessing. These methods may include removing invariant columns, removing columns based on low minor allele frequency, and some method of image-width standardization as described in the previous section. Here, however, we focus specifically on the row-sorting procedures common to these CNNs (e.g. sorting by haplotype similarity). A chosen sorting procedure has the potential to not only influence the learned decision rules, but also to restrain the power of the model to a level similar to that of previously-defined methods. If our goal is to use machine learning to build prediction methods with capabilities beyond that of current statistical approaches, then we must investigate ways to remove this human dependence.

One potential solution is to avoid the sorting step altogether, and instead implement a new machine learning architecture that is “permutation invariant,” or agnostic to the ordering of the haplotypes. To investigate the potential of this approach, we implemented a permutation-invariant CNN architecture, similar to one used recently for recombination hotspot detection [3]. Following along with previous works on permutation-invariant architectures [54], we label this model “DeepSet”. In Table 1 and S2 Table we compare the performance of the DeepSet model to the other CNNs and summary statistics. Notably, although the model under-performs in comparison to Garud’s H1 on stronger sweeps, it outperforms the other methods when applied to subtler sweeps in the simulated human populations.

While we note that DeepSet has the best performance relative to other models on very subtle sweeps (s=0.005 in the 3-population demographic model), performance is relatively low across all models and statistics in these scenarios. When we standardize images with zero-padding, DeepSet performance is artificially inflated along with that of other CNNs (see S2 Table).

## Discussion

In this study, we examine the effects of different preprocessing pipelines, CNN architectures, and simulation settings on model performance at the task of detecting selective sweeps from genome data. We then attempt to explain these models in an effort to understand how they work and when and how they might fail. An empirical analysis of the Imagene and mini-CNN models reveals that current architectures may ultimately rely on decision rules similar to that of Garud’s H1 summary statistic when row sorting is present as a preprocessing step. We hypothesize that other CNN-based sweep detection methods employing row-sorting by genetic similarity such as [14, 49, 6] act similarly. This hypothesis is supported by the findings of Flagel *et al*.[14], who note that 2-dimensional kernels provided no additional benefit over 1-dimensional kernels computing row-to-row differences. Our mini-CNN architecture, which likewise only computes differences along the rows, offers further support. In addition, to improve performance of their CNN, Deelder *et al*.[6] optimized their architecture over various hyper-parameters, finding that the optimal model contained only one convolutional layer with four filters and two dense layers. Like mini-CNN, the complexity of their architecture is strikingly small in comparison to the other works that do not discuss architecture optimization.

In this work, we observe that low complexity models (including summary statistics) seem to perform on par with the more complex models for the task of detecting selective sweeps; this suggests the possibility that the additional complexity is unnecessary in this particular context. This would be very convenient; since summary statistics like Garud’s H1 are simple to compute compared to CNN models, which require very large training sets as well as programming and machine learning expertise, this would put the power to detect sweeps into the hands of a broad swath of researchers. On the other hand, all of the models we implemented lose significant performance when selection coefficients become weaker, especially in the context of more complex demographic backgrounds. This raises the possibility that more complex models *could* detect signatures that are as-yet unobserved, given the right architecture. We also note that CNNs have been shown to be effective in other population genetics applications including identifying recombination hotspots[3] and adaptive introgression[18], and distinguishing between soft sweeps and balancing selection[23], none of which we explore here. In these cases, the additional complexity of the CNNs may well be beneficial in comparison to simpler extant models; of course, care must be taken with these due to their increased susceptibility to artifacts compared with more straightforward summary statistics.

While the particular architecture of a CNN is an important factor in its performance, an aspect of the training and testing pipeline that is perhaps equally important is also easily overlooked: the preprocessing steps used to wrangle the data into a suitable set of input images. These are necessary in part because of the peculiarities of genomic data. To represent a collection of haplotypes as an image, there are multiple decisions to be made. These include, but are not limited to: whether to use major/minor or ancestral/derived polarization for alleles at a particular site, whether to include all genome sites, or only segregating sites, whether to put a threshold on minor allele frequency for inclusion as a site, how to deal with tri-allelic sites, and the order in which to present the haplotypes. Beyond that, for a CNN classifier, all training and testing images must have uniform width, while the number of genomic sites in a simulation may be variable, depending on the choices made above.

We find that the image-width standardization preprocessing step in particular has the potential to be problematic. In the case of zero-padding, we observe a strong artifact that leads to shortcut learning by introducing a feature that is very salient to the CNN: the wider blocks of zeros required to pad sweep images relative to neutral images. As a result, the CNN performs quite well on simulated data, but would not be expected to generalize to real-world genome datasets, where the number of segregating sites in a given region is influenced not just by selection, but also by mutation and recombination rate variation. Avoiding this artifact would require the deliberate generation of training simulations based on more complex demographic models that take these factors into account. One alternative strategy of image-width standardization would be trimming images to match the minimum observed width; this would avoid the artifact associated with zero-padding, but at the expense of the losing information in the columns that are trimmed off. [Otherwise]

We are particularly interested here in the practice of row sorting, common among CNN piplelines for sweep detection, in which high-frequency haplotypes are gathered together near the top of the image. This preprocessing step appears to be a huge factor in overall performance, a finding that is supported by the work by Torada *et al*.[49], in which they find a large jump in accuracy from un-sorted to row-sorted images. Because the motivation for row-sorting is based on a signature of selection that is fairly well-understood (and serves as the basis for existing summary statistics), this raises the question of whether we are training complex models to see what we want them to see, a notion also raised by Flagel *et al*.[14]. Without this nudge, what other patterns *might* a CNN be able to find that would add to our theoretical understanding of natural selection?

There are two possible ways to address row-sorting as pushing the model in a specific direction: one can either learn an optimal sort as part of the model, or one can train a row-permutation invariant network. An immediate difficulty with the first approach is that permutations are discrete and not amenable to gradient-based training algorithms which require model operations to be continuous and differentiable. Although it has been shown that is is possible to learn a sort by way of permutation matrices[33], it is not guaranteed to be feasible or effective in this context. A distinct approach is to instead remove the sorting step entirely and focus on permutation-invariant neural networks, commonly known as deepsets [54]. By applying permutation-invariant operations on the set of individual haplotypes, these methods are entirely agnostic to the order in which the haplotypes are presented. As evidence for the potential of this approach, Chan *et al*.[3] applied a row-permutation invariant network to the task of detecting recombination rate hotspots, finding an improvement in both training speed and classification performance over a standard CNN. Furthermore, as shown by Table 1, a simple permutation invariant network outperforms the Garud’s H1 test statistic on realistic human data in the case of weak selection signals. These results provide hope for this method, and advances in architecture design and insights from other deep learning methods, such as the transformer [50], may lead to performance improvements.

Through an understanding of how high-performing machine-learning models learn patterns of evolution, we have the potential to extract new insights about these processes and the way our genomes are shaped by them. Ideally, these insights could suggest new frameworks with which to build summary statistics that can be easily constructed and deployed across a wide range of datasets. We note that this is a different perspective from which machine learning tools like CNNs are typically thought of; rather than existing as sophisticated tools only usable for completing specific tasks, we can treat them as tools that can help us learn *how* best to complete that task without the use of black-box models. We are not unique in this perspective; for instance, advances in reinforcement learning have recently lead to the development of less complex matrix multiplication algorithms discovered through the use of machine-learning models [10]. As theory concerning interpretability and explainability of machine learning improves, we can look forward to more opportunities for deriving insight into population genetics processes.

## Materials and methods

### Simulation of training and testing data

#### Simulation of a single-population demographic model

Following the “three epoch” demographic model used in Torada *et al*. [49], we generated simulations of 128 chromosomes from a single population for an 80kb region with two instantaneous population size changes: starting with an ancestral population size of 10,000, the effective population size decreases to 2,000 at 3500 generations from present, then expands to 20,000 at 3,000 generations from present. We used a mutation rate of 1.5 × 10^−8^, and a recombination rate of 1 × 10^−8^. For sweep simulations, we used selection coefficients corresponding to *s* = 0.01 for heterozygotes (0.02 for homozygotes) and *s* = 0.005 for heterozygotes (0.01 for homozygotes), with the sweep beginning 600 generations ago, and the beneficial allele located in the center of the simulated region. Simulations were generated with `msms`[8]. We hereafter refer to this as the “single-population demographic model.”

#### Simulation of 1000 Genomes populations

To simulate realistic human populations, we simulated 64 diploid individuals each from three populations representing 1000 Genomes[5] YRI, CEU, and CHB populations, across 80kb regions. Our simulations used population sizes, split times, growth rate, and migration rate parameters inferred by Gravel *et al*.[19], based on 1000 Genomes exon and low-coverage data. We used a mutation rate of 2.363 × 10^−8^ and recombination rate 1 × 10^−8^, and assumed 25 years/generation. For sweep simulations, we used a dominance coefficient of 0.5, and selection coefficients corresponding to *s* = 0.01 for heterozygotes (0.02 for homozygotes) and *s* = 0.005 for heterozygotes (0.01 for homozygotes). The beneficial allele arises 600 generations ago in the center of the 80kb region. Simulations were generated with `SLiM`[21]. We hereafter refer to this as the “three-population demographic model.”

### Preprocessing of haplotype data

For the purposes of training and testing a CNN, each set of simulated haplotypes had to be reshaped into a haplotype matrix (i.e. an image) of fixed width and length where the rows correspond to individual haplotypes and columns correspond to genome sites. Similar to the work in [49], we pre-processed the 128 sampled haplotypes from each simulation to matrix form by converting them to a binary image using major/minor polarization, removing loci with allele frequency less than 1%, standardizing the width of all images, and grouping the rows of the image into common haplotype blocks and then sorting the blocks based on haplotype frequency. To standardize the width across images, we tested two methods. First, following [49], we used the resizing algorithm from the `skimage`[52] python package to force a fixed width of 128 for each image (“image resizing”). Second, following [14], we added columns of 0s to the edges to pad the images to a specified width (“zero-padding”). In this case, for each dataset, the chosen fixed width was defined by the maximum image width rounded up to the nearest multiple of 10.

### Implementation of Deep Learning Models

The deep learning models were implemented using `Keras` [4] with `TensorFlow` [34] as the backend. Each CNN model implemented in this work is composed of a series of convolution, ReLU activation, and max pooling layers, a flattening layer, a series of dense and ReLU activation layers, and a final dense layer with a sigmoid activation function. At the start of training, the convolution and dense weights were initialized using random draws from a standard normal distribution and glorot uniform distribution [17], respectively. During training, L1 and L2 regularization terms each with a regularization penalty value of 0.005 were incorporated into the loss.

To briefly explore the potential of other deep learning architectures we also implemented a permutation invariant Deepset model [54]. The model contains two sets of convolutional and ReLU activation layers with 64 kernels of size 1×5, an averaging operation across the haplotypes, a flattening operation, 2 dense and ReLU activation layers with 64 units each, and a final dense layer with a sigmoid activation function. At the start of training, the convolution kernel and dense layer weights were initialized from the glorot uniform distribution.

For the purposes of training and testing, we partitioned each simulated dataset into a train (80%), validation (10%), and test set (10%), each with a balanced number of neutral and sweep images. Unless otherwise specified, there were 100,000 total samples for the single-population model (generated with `msms`), and 20,000 total samples in the three-population model (generated with `SLiM`) (see S10 Fig for sample complexity analysis). The ML models were trained to distinguish between selective sweep and neutral images by minimizing the standard binary cross-entropy loss function. The Adam optimizer [25] with a mini batch size of 64, settings *β*_1_ = 0.9 and *β*_2_ = 0.999, and learning rate of 0.001 was used. All models were trained for 2 epochs where 1 epoch is a single pass through the training dataset. Each model was trained 10 times on the training set, and the best performing model on the validation set was used for testing and model comparison. To test the performance of each model, accuracy, area under the ROC curve, and correlation estimates were calculated using the balanced test set. All models were trained on a combination of Nvidia RTX 2060 and 3090 GPUs.

### Implementation of summary statistics

Each summary statistic was implemented and applied to the raw haplotype data using the `scikit-allel` python package [35]. iHS values were standardized based on neutral simulations for the demographic model of interest, learning the mean and variance for standardization for each derived allele count from 0 to 128. To produce a single value for the whole genome region, we took the maximum absolute value of all standardized iHS values across the region.

For each summary statistic, a decision threshold was used to classify a region as sweep or neutral based on the computed statistic. This threshold was found by numerically computing the optimal decision threshold which maximized the accuracy of the method on the training set, and performance was evaluated on the test set.

### Methods of interpretation

Where possible, we plotted visualizations of the convolution kernel and dense weights. In the case of more complex models with large parameter counts, we used the SHAP DeepExplainer [32] to generate visual explanations for model predictions. For a given deep learning model and input, SHAP values are used to measure the importance of input features to the final prediction. When visualized, the red pixels may be interpreted as feature locations contributing to an increase in the model’s output (towards sweep classification) while blue pixels contribute to a decrease (towards neutral classification). For each simulation type, we provided 20 neural and 20 sweep images for use by DeepExplainer as background samples.

## Supporting information

**S1 File. Convolutional Neural Networks for Selection Detection in the Literature**.

**S1 Fig. Comparison of Imagene with mini-CNN**.

**S1 Table. Model accuracy for decreasing levels of complexity, for single-population demographic model with selection coefficient** *s* = 0.01.

**S2 Table. Model accuracy for all CNN models and summary statistics, across all demographic models and types of pre-processing**.

**S2 Fig. Performance correlation for summary statistics and CNN approaches under image resizing**.

**S3 Fig. Performance correlation for summary statistics and CNN methods under zero-padding**.

**S4 Fig. Simulations of CEU and YRI under the three-population demographic model match the site frequency spectrum of 1000 Genomes populations**.

**S5 Fig. SHAP explanations for Imagene predictions under image resizing**.

**S6 Fig. SHAP explanations for Imagene predictions under zero-padding**.

**S7 Fig. Visualization of mini-CNN dense layer under image resizing**.

**S8 Fig Visualization of mini-CNN dense layer under zero-padding**.

**S9 Fig. Full images corresponding to the cropped images in Figure 4**.

**S10 Fig. Sample Complexity Analysis**.

## Acknowledgments

We are grateful to Sara Mathieson, Arthur Sugden, and members of the Ramachandran lab for valuable discussion and feedback. We are also grateful to Matteo Fumagalli for sharing resources related to Imagene.

## Software Availability

All code for reproducing the results in this paper is available at https://github.com/ryanmcecil/popgen_ml_sweep_detection.

## References

[1] Tom Brown et al. “Language Models are Few-Shot Learners”. In: Advances in Neural Information Processing Systems. Ed. by H. Larochelle et al. Vol. 33. Curran Associates, Inc., 2020, pp. 1877–1901. URL: https://proceedings.neurips.cc/paper/2020/file/1457c0d6bfcb4967418bfb8ac142f64a-Paper.pdf.

[2] Jedidiah Carlson et al. “Extremely rare variants reveal patterns of germline mutation rate heterogeneity in humans”. In: Nature communications 9.1 (2018), pp. 1–13.

[3] Jeffrey Chan et al. “A Likelihood-Free Inference Framework for Population Genetic Data using Exchangeable Neural Networks”. In: bioRxiv (2018). doi: 10.1101/267211. eprint: https://www.biorxiv.org/content/early/2018/11/05/267211.full.pdf. URL: https://www.biorxiv.org/content/early/2018/11/05/267211.

[4] François Chollet et al. Keras. https://keras.io. 2015.

[5] 1000 Genomes Project Consortium et al. “A global reference for human genetic variation”. In: Nature 526.7571 (2015), p. 68.

[6] Wouter Deelder et al. “Using deep learning to identify recent positive selection in malaria parasite sequence data”. In: Malar J 20 (2021). doi: https://doi.org/10.1186/s12936-021-03788-x. URL: https://malariajournal.biomedcentral.com/track/pdf/10.1186/s12936-021-03788-x.pdf.

[7] Jacob Devlin et al. “BERT: Pre-training of Deep Bidirectional Transformers for Language Understanding”. In: Proceedings of the 2019 Conference of the North American Chapter of the Association for Computational Linguistics: Human Language Technologies, Volume 1 (Long and Short Papers). Minneapolis, Minnesota: Association for Computational Linguistics, June 2019, pp. 4171–4186. doi: 10.18653/v1/N19-1423. URL: https://aclanthology.org/N19-1423.

[8] Gregory Ewing and Joachim Hermisson. “MSMS: a coalescent simulation program including recombination, demographic structure and selection at a single locus”. In: Bioinformatics (Oxford, England) 16 (2010). doi: 10.1093/bioinformatics/btq322.

[9] Arnaud Nguembang Fadja et al. “Identification of natural selection in genomic data with deep convolutional neural network”. In: BioData Mining 14 (2021). doi: https://doi.org/10.1186/s13040-021-00280-9. URL: https://biodatamining.biomedcentral.com/track/pdf/10.1186/s13040-021-00280-9.pdf.

[10] Alhussein Fawzi et al. “Discovering faster matrix multiplication algorithms with reinforcement learning”. In: Nature 610.7930 (2022), pp. 47–53.

[11] Justin C Fay and Chung-I Wu. “Hitchhiking under positive Darwinian selection”. In: Genetics 155.3 (2000), pp. 1405–1413.

[12] Anna Ferrer-Admetlla et al. “On detecting incomplete soft or hard selective sweeps using haplotype structure”. In: Molecular biology and evolution 31.5 (2014), pp. 1275–1291.

[13] Anna Ferrer-Admetlla et al. “On detecting incomplete soft or hard selective sweeps using haplotype structure”. In: Molecular biology and evolution 31.5 (2014), pp. 1275–1291.

[14] Lex Flagel, Yaniv Brandvain, and Daniel R Schrider. “The Unreasonable Effectiveness of Convolutional Neural Networks in Population Genetic Inference”. In: Molecular Biology and Evolution 36.2 (Dec. 2018), pp. 220–238. ISSN: 0737-4038. doi: 10.1093/molbev/msy224. eprint: https://academic.oup.com/mbe/article-pdf/36/2/220/27736968/msy224.pdf. URL: https://doi.org/10.1093/molbev/msy224.

[15] Nandita R. Garud et al. “Recent selective sweeps in north american drosophila melanogaster show signatures of soft sweeps”. In: PLoS Genetics (2015). doi: 10.1371/journal.pgen.1005004.

[16] Robert Geirhos et al. “Shortcut learning in deep neural networks”. In: Nature Machine Intelligence 2.11 (2020), pp. 665–673.

[17] Xavier Glorot and Yoshua Bengio. “Understanding the difficulty of training deep feedforward neural networks”. In: Proceedings of the thirteenth international conference on artificial intelligence and statistics. JMLR Workshop and Conference Proceedings. 2010, pp. 249–256.

[18] Graham Gower et al. “Detecting adaptive introgression in human evolution using convolutional neural networks”. In: Elife 10 (2021), e64669.

[19] Simon Gravel et al. “Demographic history and rare allele sharing among human populations”. In: Proceedings of the National Academy of Sciences 108.29 (2011), pp. 11983–11988.

[20] Sharon R Grossman et al. “A composite of multiple signals distinguishes causal variants in regions of positive selection”. In: Science 327.5967 (2010), pp. 883–886.

[21] Benjamin C Haller and Philipp W Messer. “SLiM 3: Forward Genetic Simulations Beyond the Wright–Fisher Model”. In: Molecular Biology and Evolution 36.3 (Jan. 2019), pp. 632–637. ISSN: 0737-4038. doi: 10.1093/molbev/msy228. eprint: https://academic.oup.com/mbe/article-pdf/36/3/632/27980602/msy228.pdf. URL: https://doi.org/10.1093/molbev/msy228.

[22] Kaiming He et al. “Deep Residual Learning for Image Recognition”. In: 2016 IEEE Conference on Computer Vision and Pattern Recognition (CVPR). 2016, pp. 770–778. doi: 10.1109/CVPR.2016.90.

[23] Ulas Isildak, Alessandro Stella, and Matteo Fumagalli. “Distinguishing between recent balancing selection and incomplete sweep using deep neural networks”. In: Molecular Ecology Resources 21 (2021). doi: https://doi.org/10.1111/1755-0998.13379. URL: https://onlinelibrary.wiley.com/doi/epdf/10.1111/1755-0998.13379.

[24] John Jumper et al. “Highly accurate protein structure prediction with AlphaFold”. en. In: Nature 596.7873 (Aug. 2021), pp. 583–589.

[25] Diederik P. Kingma and Jimmy Ba. “Adam: A Method for Stochastic Optimization”. In: 3rd International Conference on Learning Representations, ICLR 2015, San Diego, CA, USA, May 7-9, 2015, Conference Track Proceedings. Ed. by Yoshua Bengio and Yann LeCun. 2015. URL: http://arxiv.org/abs/1412.6980.

[26] Erich Kobler et al. “Total Deep Variation for Linear Inverse Problems”. In: IEEE Conference on Computer Vision and Pattern Recognition. 2020.

[27] Augustine Kong et al. “A high-resolution recombination map of the human genome”. In: Nature genetics 31.3 (2002), pp. 241–247.

[28] Alex Krizhevsky, Ilya Sutskever, and Geoffrey E Hinton. “ImageNet Classification with Deep Convolutional Neural Networks”. In: Advances in Neural Information Processing Systems. Ed. by F. Pereira et al. Vol. 25. Curran Associates, Inc., 2012. URL: https://proceedings.neurips.cc/paper/2012/file/c399862d3b9d6b76c8436e924a68c45b-Paper.pdf.

[29] Y. Lecun et al. “Gradient-based learning applied to document recognition”. In: Proceedings of the IEEE 86.11 (1998), pp. 2278–2324. doi: 10.1109/5.726791.

[30] Kao Lin et al. “Distinguishing positive selection from neutral evolution: boosting the performance of summary statistics”. In: Genetics 187.1 (2011), pp. 229–244.

[31] Pantelis Linardatos, Vasilis Papastefanopoulos, and Sotiris Kotsiantis. “Explainable AI: A review of machine learning interpretability methods”. en. In: Entropy (Basel) 23.1 (Dec. 2020), p. 18.

[32] Scott M Lundberg and Su-In Lee. “A Unified Approach to Interpreting Model Predictions”. In: Advances in Neural Information Processing Systems 30. Ed. by I. Guyon et al. Curran Associates, Inc., 2017, pp. 4765–4774. URL: http://papers.nips.cc/paper/7062-a-unified-approach-to-interpreting-model-predictions.pdf.

[33] Jiancheng Lyu et al. “AutoShuffleNet: Learning Permutation Matrices via an Exact Lipschitz Continuous Penalty in Deep Convolutional Neural Networks”. In: Proceedings of the 26th ACM SIGKDD International Conference on Knowledge Discovery Data Mining. KDD ‘20. Virtual Event, CA, USA: Association for Computing Machinery, 2020, pp. 608–616. ISBN: 9781450379984. doi: 10.1145/3394486.3403103. URL: https://doi.org/10.1145/3394486.3403103.

[34] Martín Abadi et al. TensorFlow: Large-Scale Machine Learning on Heterogeneous Systems. Software available from http://tensorflow.org. 2015. URL: https://www.tensorflow.org/.

[35] Alistair Miles et al. cggh/scikit-allel: v1.3.3. 2021.

[36] Volodymyr Mnih et al. Playing Atari with Deep Reinforcement Learning. 2013. doi: 10.48550/ARXIV.1312.5602. URL: https://arxiv.org/abs/1312.5602.

[37] Marc Pybus et al. “Hierarchical boosting: a machine-learning framework to detect and classify hard selective sweeps in human populations”. In: Bioinformatics 31.24 (2015), pp. 3946–3952.

[38] Gabrielle Ras et al. “Explainable deep learning: A field guide for the uninitiated”. In: J. Artif. Intell. Res. 73 (Jan. 2022), pp. 329–397.

[39] Cynthia Rudin et al. “Interpretable machine learning: Fundamental principles and 10 grand challenges”. In: Statistics Surveys 16.none (2022), pp. 1–85. doi: 10.1214/21-SS133. URL: https://doi.org/10.1214/21-SS133.

[40] Pardis C Sabeti et al. “Detecting recent positive selection in the human genome from haplotype structure”. In: Nature 419.6909 (2002), pp. 832–837.

[41] Pardis C Sabeti et al. “Genome-wide detection and characterization of positive selection in human populations”. In: Nature 449.7164 (2007), pp. 913–918.

[42] Daniel R Schrider and Andrew D Kern. “S/HIC: robust identification of soft and hard sweeps using machine learning”. In: PLoS genetics 12.3 (2016), e1005928.

[43] Daniel R Schrider and Andrew D Kern. “Supervised machine learning for population genetics: a new paradigm”. In: Trends in Genetics 34.4 (2018), pp. 301–312.

[44] Andrew W Senior et al. “Improved protein structure prediction using potentials from deep learning”. en. In: Nature 577.7792 (Jan. 2020), pp. 706–710.

[45] Sara Sheehan and Yun S Song. “Deep learning for population genetic inference”. In: PLoS computational biology 12.3 (2016), e1004845.

[46] David Silver et al. “Mastering the game of Go without human knowledge”. en. In: Nature 550.7676 (Oct. 2017), pp. 354–359.

[47] Lauren Alpert Sugden et al. “Localization of adaptive variants in human genomes using averaged one-dependence estimation”. In: Nature communications 9.1 (2018), pp. 1–14.

[48] Fumio Tajima. “Statistical method for testing the neutral mutation hypothesis by DNA polymorphism.” In: Genetics 123.3 (1989), pp. 585–595.

[49] Luis Torada et al. “ImaGene: a convolutional neural network to quantify natural selection from genomic data”. In: BMC Bioinformatics 20 (2019). doi: https://doi.org/10.1186/s12859-019-2927-x. URL: https://bmcbioinformatics.biomedcentral.com/articles/10.1186/s12859-019-2927-x.

[50] Ashish Vaswani et al. “Attention is All you Need”. In: Advances in Neural Information Processing Systems. Ed. by I. Guyon et al. Vol. 30. Curran Associates, Inc., 2017. URL: https://proceedings.neurips.cc/paper/2017/file/3f5ee243547dee91fbd053c1c4a845aa-Paper.pdf.

[51] Benjamin F Voight et al. “A map of recent positive selection in the human genome”. In: PLoS biology 4.3 (2006), e72.

[52] Stéfan van der Walt et al. “scikit-image: image processing in Python”. In: PeerJ 2 (June 2014), e453. ISSN: 2167-8359. doi: 10.7717/peerj.453. URL: https://doi.org/10.7717/peerj.453.

[53] Tom Young et al. “Recent Trends in Deep Learning Based Natural Language Processing [Review Article]”. In: IEEE Computational Intelligence Magazine 13.3 (2018), pp. 55–75. doi: 10.1109/MCI.2018.2840738.

[54] Manzil Zaheer et al. “Deep Sets”. In: Advances in Neural Information Processing Systems. Ed. by I. Guyon et al. Vol. 30. Curran Associates, Inc., 2017. URL: https://proceedings.neurips.cc/paper/2017/file/f22e4747da1aa27e363d86d40ff442fe-Paper.pdf.

